# Impact of storage on starch digestibility and texture of a high-amylose wheat bread

**DOI:** 10.1101/2022.07.04.498686

**Authors:** M. Corrado, P. Zafeiriou, J.H. Ahn-Jarvis, G.M. Savva, C.H. Edwards, B.A. Hazard

**Affiliations:** Food Innovation and Health, Quadram Institute Bioscience, Norwich Research Park, UK; Designing Future Wheat and Molecules from Nature, John Innes Centre, Norwich Research Park, UK

**Keywords:** amylolysis, starch branching enzyme II, chain-length distribution, birefringence, staling, bread, firming, digestion

## Abstract

Staling is a complex process that determines the shelf-life of baked products like bread. Breads made using high-amylose flour may elicit a lower glycaemic response, with benefits for health, however the impact of storage on novel high-amylose wheat foods structure are not known.

We investigated the staling behaviour of high-amylose bread made from a *starch branching enzyme II* (*sbeII*) wheat mutant compared to a wild-type (WT) control, by measuring starch digestibility (susceptibility to amylolysis) and bread texture over time in different storage conditions. Breads prepared from *sbeII* and WT control wheat flours were subjected to fresh, refrigerated and frozen storage, and starch digestibility and crumb texture were measured up to three days. Starch from *sbeII* flour was characterised by a larger proportion of long chains resulting in increased amylose content, typical of *sbeII* mutant wheat. Starch in *sbeII* bread was less susceptible to amylolysis when freshly baked (~17% difference of starch digested at 90 min, C_*90*_) and after storage (26%-28% C_*90*_ difference, depending on the storage condition), compared to the WT control. Texture of freshly baked *sbeII* bread was similar to the WT control; storage conditions affected the progression of crumb firming and resilience to touch for both breads, but changes in crumb texture were less pronounced in *sbeII* bread. Overall, *sbeII* bread was less prone to staling than conventional WT bread during the first three days of storage, particularly when stored in the fridge or at room temperature.

## Introduction

Bread made from white wheat flour is a staple food in many countries, however, its high glycaemic potency can potentially harm cardiometabolic health over time (Livesey, et al., 2019).

The glycaemic response to bread reflects the digestibility of starch, the main dietary carbohydrate, so developing wheat starch with greater resistance to digestive enzymes (such as pancreatic α-amylase) may lead to a viable healthier alternative to conventional white wheat bread, if product quality and shelf-life can be preserved.

One promising approach is the use of novel types of wheat with increased amylose content and lower starch digestibility (Hallström E., Sestili F., Lafiandra D., Björck I., & Östman E., 2011; Roman L. & Martinez M. M., 2019). *Starch branching enzyme II* (*sbeII*) wheat mutants obtained by Targeting Induced Local Lesions in Genomes (TILLING), a non-transgenic technology (McCallum, Comai, Greene, & Henikoff, 2000), can produce starch with higher amylose content than conventional wheat starch (Schonhofen A., Zhang X., & Dubcovsky J., 2017). Foods made from *sbeII* wheat flour have lower starch digestibility and could therefore be used to lower glycaemic responses to starch-rich staple foods (Corrado M., et al., 2022; Corrado M., et al., 2020; Sissons M., Sestili F., Botticella E., Masci S., & Lafiandra D., 2020).

The use of high-amylose foods to replace conventional high glycaemic ones requires studies of their ageing behaviour, a determinant of shelf-life. Starch structure determines its physicochemical properties during storage, which underpin not only digestibility but also bread texture and quality.

The bread making process affects both the starch and protein fractions in the flour while hydration, shear, and heat determine gluten development and starch gelatinisation. During storage, moisture loss and starch retrogradation and gluten network changes contribute to staling causing increased hardness over time. This is due to recrystallization of the starch fraction (Ribotta P.D., Leon A.E., & Anon M.C., 2003), water migration from crumb to crust and between protein and starch favouring network plasticisation and increased stiffness (Bosmans G. M., Lagrain B., Fierens E., & Delcour J. A., 2013). Starch, of which 20%-30% is amylose and 70%-80% is amylopectin in wheat, can retrograde by forming new ordered structures within the gelatinised granule (Miles M. J., Morris V. J., Orford P. D., & Ring S. G., 1985). The molecular structure of these two polymers determines starch physicochemical characteristics and functionality during processing (Vamadevan & Bertoft, 2018). Considering starch (re-)crystallisation during storage, differences in the molecular structure affect this dynamic process greatly. Amylose is characterised by infrequent branches and can crystallise quickly, while amylopectin, a highly branched polymer, requires more time to form new double helical structures (Kohyama K., Matsuki J., Yasui T., & Sasaki T., 2004; Roman L., Campanella O., & Martinez, 2019).

Bread ageing (staling) and consequently shelf-life are determined by texture firming (network plasticization), moisture migration and starch retrogradation (Gray J.A. & Bemiller J.N., 2003). Changes in the molecular structure of starch, the major component of wheat flour, can affect starch physicochemical properties and the progression of staling depending on the degree of the modification. For instance, Arp *et al*. showed that 10% and 20% flour replacement with high amylose maize starch (Hi-Maize 260™) in bread led to similar firming behaviour as the control while 30% replacement led to higher firmness (Arp C.G., Correa M.J., & Ferrero C., 2020). Only few studies have investigated the functionality of high amylose starch in bread obtained by starch structural modification *in planta*. In two recent studies, bread made from high-amylose wheat flour (51.5%) had similar texture to the control bread (Li C., Dhital S., & Gidley M. J., 2022). During fridge storage, high amylose bread was found to be less prone to ageing than the control bread (Li C. & Gidley M.J., 2022). Improving the understanding of the physicochemical transformations of high-amylose *sbeII* wheat starch during baking and storage are of interest as they can affect texture and ultimately, palatability and shelf-life of this food product.

In this study, we investigated the effect of different conditions commonly used to store bread on moisture loss, crumb texture and starch susceptibility to amylolysis in a *sbeII* bread compared to a wild-type (WT) control bread. Breads were produced using a modified straight dough method which resembles a traditional home-baking process and analysed freshly baked (after 2h of cooling corresponding to the moment breads can be packaged and stored, later referred to as ‘0h’) and after storage at room temperature (RT), in the fridge and in the freezer for up to 72 hours as at lower storage temperature starch is expected to retrograde faster with evident changes in crumb firmness (Bosmans G. M., Lagrain B., Fierens E., et al., 2013). Structural differences between *sbeII* mutant starch and the WT control were measured using size exclusion chromatography (SEC) and starch crystallinity differences between *sbeII* and WT control flour and breads were explored using light microscopy, in raw flour, in freshly baked bread to identify differences due to the baking process and after freezer storage (7 days) as indicator of bread ageing.

## Materials and methods

### Wheat materials and bread making process

A field trial at the John Innes Centre Field Station (Norfolk, UK) was used to supply wheat grains of a high-amylose *sbeII* mutant and WT control (Schonhofen A., et al., 2017). Grains were milled to produce refined white flour which was used for making bread rolls with approximately 75 g of total starch per roll. The starch characteristics of the flour used in this study have been described previously (Corrado M., et al., 2022). Briefly, the total starch content of the *sbeII* and WT control flours was similar while, as expected, the apparent amylose proportion measured by iodine-binding method on starch isolated from flour was greater in *sbeII* starch (39 ± 1.1% of total starch, mean ± SEM, n = 3) than the WT control (25.7 ± 0.7% of total starch, mean ± SEM, n = 3).

Bread rolls were baked from *sbeII* and WT flours as described previously (Corrado M., et al., 2022) to achieve a similar starch content (~75 g) based on the total starch content of the flour. The same process as used to produce *sbeII* and WT control doughs, later portioned into rolls and baked. Using a Kenwood mixer (KM300, Kenwood UK) with a hook attachment, flour (WT = 57%, *sbeII* = 55%), caster sugar (WT and *sbeII* = 2%), Allison’s dry yeast (WT and *sbeII* = 2%) and water (WT = 37%, *sbeII* = 39%) were mixed at low speed for one minute, after which Willow vegetable shortening (WT and *sbeII* = 2%) was added. After three minutes of mixing, salt (WT and *sbeII* = 1%) was added and the dough was mixed at increasing speed for five more minutes.

The dough was then fermented for two hours at ambient conditions (21°C, 41% Relative Humidity (RH)), then rolls were shaped and proofed for 15 minutes at 40°C, 100% RH using a steam oven (DG 6001 GourmetStar, Miele, UK). Rolls were sprayed with water and baked at ~185°C for 20 minutes in a pre-heated convection oven (Hotpoint HAE60P, Whirlpool UK) with a tray of water to provide steam, to reach a core temperature of ~95°C. Batches were prepared under identical conditions and rolls (3 per batch) were paired by their baking position in the oven to ensure even baking. After baking, rolls were left at room temperature to cool for two hours. Proximate nutritional content of breads is reported in Supplementary Table 1.

### Allocation to storage conditions and sample preparation

After 2h cooling at room temperature, bread rolls were either analysed immediately (fresh, 0h) or after 24h, 48h, and 72h of storage at room temperature (RT, +19 to +21°C), freezer temperature (−18 to −20°C) and fridge temperature (+3 to +5°C), n = 3 per condition. Samples stored in the fridge and freezer were allowed to return to RT before analysis. A roll of ~165 g required approximately 3h to reach RT at the core after freezer storage, and approximately 2h after fridge storage.

Figure 1 shows sample preparation of each bread genotypes for the different analyses.

**Figure 1.**
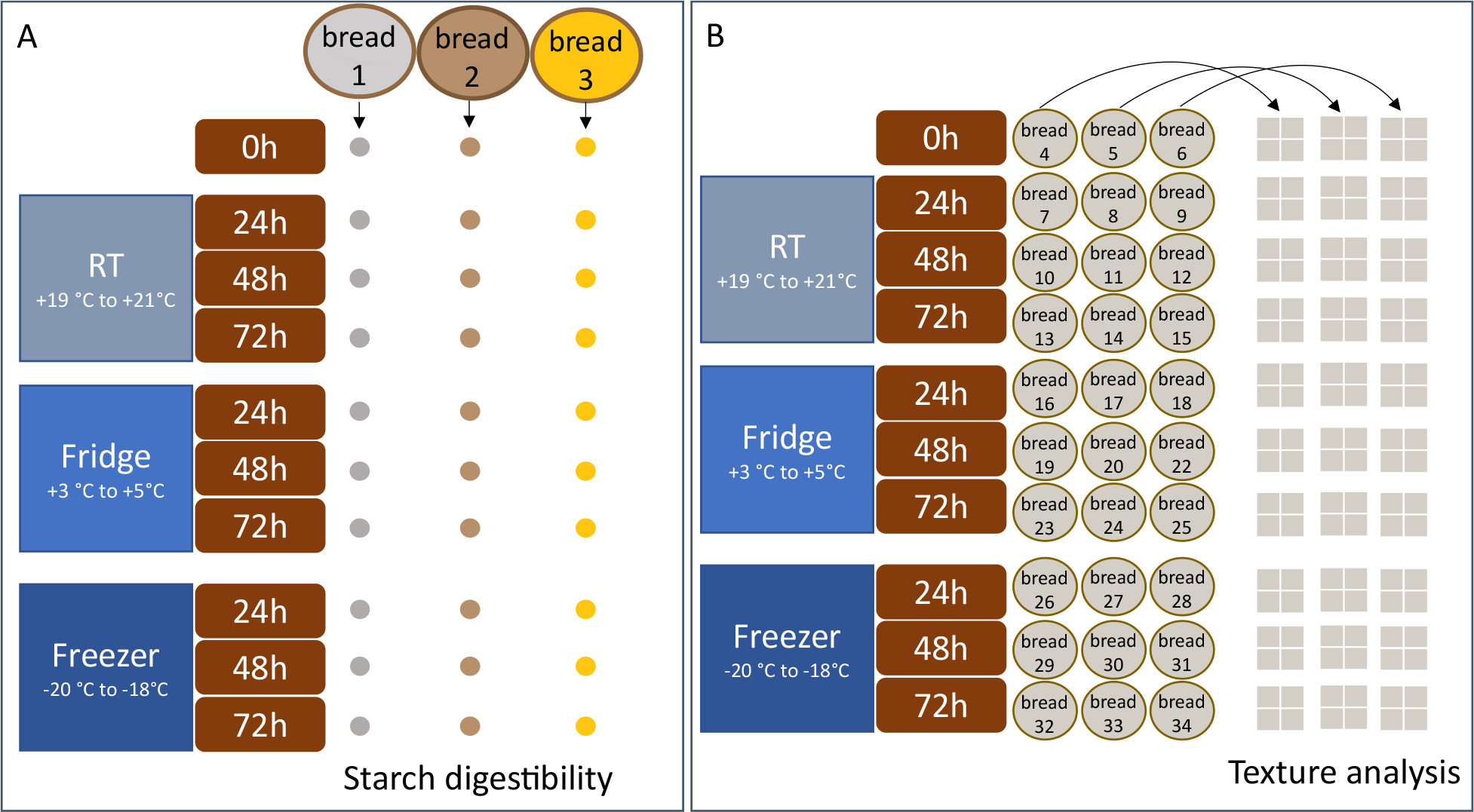
Bread sampling for starch digestibility (A), texture analysis (B). For starch digestibility, seven samples per genotype (sbeII and WT) were taken from each bread roll (●) after baking and allocated to a storage condition (n = 3 per storage condition). Isolated crumb from each roll was ground and sieved using 1 mm sieve. The fraction below 1 mm was used for in vitro starch susceptibility to amylolysis; the fraction above 1 mm was used for moisture analysis. Samples for the starch digestibility assay and moisture analysis were weighed out at the same time and either analysed or stored at the storage conditions described above. For texture analysis, three rolls per genotype (sbeII and WT) were assigned to each storage condition. For the analysis, the crust of the bread was removed to isolate the core crumb which was sliced into four 5×5×5 cm cubes (□) and used for texture analysis (TPA).

**For starch digestibility analysis**, the structure of the bread was not preserved as the starch characteristics were analysed at microstructural level.

Three bread rolls per bread type (*sbeII* and WT control) were produced, with a sample from each roll subjected to each storage condition. After 2h of cooling at RT, the crust was removed and the crumb was ground using a Kenwood mini-chopper food processor and sieved using a 1 mm sieve. The fraction below 1 mm was used to determine starch digestibility *in vitro*. Starch susceptibility to amylolysis was measured on fresh bread and bread stored for 24h, 48h, 72h at RT, fridge and freezer conditions, as described above.

All samples were weighed out from fresh bread and then stored in the allocated conditions, with three independent samples (from three rolls) per condition, Figure 1A. For samples stored in cold temperature (fridge and freezer), tubes were allowed to return to RT before analysis. Because of the small amount of sample required for this type of analysis (~63 mg) compared to the weight of a whole roll (~165 g), thawing time was proportionally adjusted. Samples from the fridge reached RT in 20 min and samples from the freezer reached RT in 45 minutes.

**For texture and moisture analyses**, the macrostructure of the rolls was preserved as the crust plays a role in moisture loss and crumb firming. Therefore, bread rolls were produced over three subsequent days and assigned to a storage condition (n = 3 rolls per condition). Each roll was stored in fridge, freezer or at RT in individual sealed bags.

Each analysis was carried out on three independent rolls, from different batches of dough to capture batch-to-batch variability and ensure fair comparison between *sbeII* and WT control breads. After storage and ahead of analysis, each roll was weighed then the crust was discarded, the core of the roll was cut into four 5×5×5 cm cubes and was analysed immediately (leading to four technical replicates for each measure), Figure 1B. Moisture by air-oven method AACC (44-15A), one stage procedure (AACC International, 1999) was used to measure moisture in samples from the same rolls used for texture analysis, using the remaining bread crumb (n = 3 independent samples per condition).

**For microscopy**, three rolls were produced per bread type (*sbeII* and WT control), samples of crust and crumb (from the core of the roll) were taken when fresh (2h after baking) after which the rolls were stored in the freezer for 7 days. After 7 days, each roll was left at RT for approx. 3h to reach ambient temperature (+19°C). After thawing, crust and crumb samples were taken from the core of the roll and mixed with water to disrupt the crumb matrix and imaged.

## Analytical Methods

### Moisture of bread crumb during storage

Moisture content of bread crumb samples was measured by the air-oven drying (AACC 44-15A), one stage procedure (AACC International, 1999). Samples were weighed out in metal tins immediately after slicing to prevent moisture loss. These were then placed in the oven for exactly 16 hours, then removed from the oven and left to cool for one hour in a desiccator before weighing the tins again. Moisture content was calculated by subtracting the weight of the sample after drying in the oven to that of the fresh sample for three independent replicates (one from each roll).

Moisture content of bread rolls during storage was also determined by weighing the rolls before and after storage.

### Texture analysis of bread crumb during storage

Crumb texture was determined using a two-bite test to obtain *Hardness*, *Cohesiveness*, *Chewiness*, *Gumminess*, *Springiness*, and *Resilience* of bread on a TA-XT2 Texture Analyser (Stable Micro System, Godalming, UK) equipped with a 5-kg load cell and 50 mm compression plate (P50). Uniaxial compression with 100 mm min-1 crosshead speed was applied to a 5 × 5 × 5 cm sample with crumb hardness corresponding to the force *N* required for a 40% compression. Parameters of interest were obtained using Exponent software (6.0, Stable Micro System, Godalming, UK), (Corrado M., et al., 2022).

### Starch susceptibility to amylolysis

Starch susceptibility to amylolysis was measured on bread crumb. Sieved bread crumb was weighed in a tube to achieve 27 mg of starch in 5 mL of Phosphate-Buffered Saline (PBS). Samples were incubated at 37°C with end-over mixing and 2U/mL of porcine pancreatic α-amylase per incubation mix were added to start the assay, after taking a baseline measurement of endogenous maltose (Y0). Samples of incubations mix (100 μl) were taken at 0 (before adding the enzyme), 3, 6, 9, 12, 15, 18, 21, 25, 30, 35, 45, 60, 75, 90 minutes of incubation and added to tubes containing 100 μl of Na_2_CO_3_ to stop the reaction. Reducing sugars obtained from hydrolysis were measured using ‘PAHBAH’ (p-hydroxybenzoic acid hydrazide) colorimetric method, (Edwards C.H., Cochetel N., Setterfield L., Perez-Moral N., & Warren F.J., 2019). Samples were centrifuged 15,000 × g for 5 minutes; the supernatant was diluted 1:10 with deionised water to 100 μL and incubated in a boiling waterbath for 5 minutes with 1 mL of PAHBAH reagent. For the standard curve, 1 mM maltose in water was used to make up the standard solutions (0 to 1 mM) and incubated with PAHBAH reagent with the digestion samples.

Absorbance was read in a microplate reader (VersaMax Microplate Reader, Molecular Devices, LLC., CA, USA) at 405 nm and maltose equivalent concentration in samples were calculated using the maltose standard curve.

### Starch chain-length distribution

Starch isolated from *sbeII* and WT wheat flour was used to determine starch chain length distribution by size exclusion chromatography (SEC). For the starch isolation, ~5g of flour were mixed with 30 mL of deionised water then filtered through a layer of Miracloth (475855, Calbiochem®). The solution was centrifuged (2000 × g for 5 min) and the recovered pellet was washed once with 30 mL of 2% Sodium Dodecyl Sulfate (SDS) and then centrifuged. This process was repeated twice using 5 mL of 2% SDS. The resulting starch pellet was washed three times with 5 mL of acetone and left to dry overnight to ensure complete evaporation of the solvent (Corrado M., et al., 2020; Verhoeven, Fahy, Leggett, Moates, & Denyer, 2004). Starch (8 mg/mL) was solubilised in DMSO containing 0.5% (w/w) LiBr followed by enzymatic debranching with isoamylase (Wu, Li, & Gilbert, 2014), freeze-dried, and analysed by HPLC-SEC. Samples and standards were resolubilised in 4 and 2 mg/mL, respectively with DMSO+0.5% (w/w), and stored at 80°C overnight prior to analysis. The HPLC-SEC system for peak resolution was a Waters Alliance e2695 HPLC (Milford, MA, USA) equipped with refractive index detector (RI), autosampler (40°C), and column heater (90°C), as described by (Fahy B., Gonzalez O., Savva G. M., et al., 2022).

The isocratic mobile phase was prepared with DMSO with 0.5% LiBr (w/w) at 0.500 mL/min flow rate. Two analytical columns (10 um; 300 Å followed by 30 Å; 8 × 300 mm, GRAM; Polymer Standard Service, Mainz, DE) with a guard column (8 × 50 mm, GRAM; Polymer Standard Service, Mainz, DE) were used as stationary phase. The total run time was 65 minutes and injection volume for each sample was 50 μL. Calibration curves were generated using pullulan standards (PSS-pulkit, Polymer Standard Service, Mainz, DE) with peak molecular weights ranging from 342 to 708,000 Da and correlation coefficients of R2 = 0.9993 ± 0.0005. The relationship between elution volume and hydrodynamic radius (Vh) for the linear glucans was determined using calibration curves described similarly by Cave, *et al*.(Cave, Seabrook, Gidley, & Gilbert, 2009). The debranched starch samples RI elution profiles were converted to SEC weight distributions as described in detail by Perez-Moral, *et al*. (Perez-Moral, Plankeele, Domoney, & Warren, 2018). The relative peak heights corresponding to amylopectin and amylose chains were calculated as defined by Hanashiro *et al*. 1996 (Hanashiro, Abe, & Hizukuri, 1996) and Vilaplana *et al*. 2012 (Vilaplana, Hasjim, & Gilbert, 2012).

### Microscopy

Images of starch were obtained using an Olympus BX60 upright brightfield and widefield fluorescent microscope with colour camera, using a brightfield and polarised light filter to visualise the birefringence pattern, objective ×10. Images were captured using the ProgRes CT3 Colour 3.15 MP cMOS camera: 3.2 μm2 pixel size, 2048 × 1536 image size.

Samples of crust and crumb were mixed with water briefly to disrupt the bread matrix. The suspension was laid on microscope slides 1.0 – 1.2 mm with glass cover n. 1.5, 22 × 50mm, (SuperFrost®, VWR®) and imaged immediately.

### Statistical analysis

Amylolysis curves were fitted to a first order equation previously described (Edwards C.H., et al., 2019) using a non-linear regression model. The first order rate constant (*k*) and predicted starch digested at the end of reaction (*C_∞_)* were estimated from the equation, after subtracting the ‘endogenous’ maltose detected before the start of the reaction (Y_0_) from the subsequent timepoints. The experimental endpoint *C_90_* and the incremental Area Under the Curve (iAUC) are reported as additional descriptors of the susceptibility to hydrolysis.

Amylolysis parameters (*C_90_* and iAUC, *k* and *C_∞_*) were compared across groups with mixed-effects models using the *lmerTest* R package (version 3.1.2), (Bates, Mächler, Zurich, Bolker, & Walker, 2014; Kuznetsova A., Brockhoff P.B., & Christensen R.H.B., 2017) with storage and genotype as fixed effects and an interaction term of storage condition by genotype.

A separate mixed-effects model was estimated for each texture parameter following log-transformation. Upon visual inspection, log-transformation stabilised the variances and resulted in model residuals normally distributed. Each model included fixed effects of every combination of storage condition (storage duration by storage temperature), the main effect of genotype and the interaction between storage condition and genotype as well as a random intercept to account for having multiple data points from each roll. With regard to springiness, data was difficult to model because of variability between replicates, so no statistical model was fitted for this outcome (Supplementary Figure 2).

The effect of storage on bread moisture was estimated using a linear regression model, with main effects of storage condition and genotype and an interaction term of storage condition by genotype.

Chain-length distribution of debranched starch was analysed as described by Fahy and colleagues (Fahy B., Gonzalez O., Savva G.A., et al., 2022) using a mixture model where the distribution components were estimated using constrained values of degree of polymerisation (DP) previously described by Hanashiro *et al*. 1996 (Hanashiro, et al., 1996) and Vilaplana *et al*. 2012 (Vilaplana, et al., 2012). The *mixdist* (version 0.5-5) R package was used to estimate mixture models (Macdonald P. & Du J., 2018).

Specific contrasts and marginal means were obtained from mixed models with Satterthwaite approximations for degrees of freedom, using the *emmeans* R package (version1.7.2), following back-transformation, if required (Lenth R., 2020).

Datasets were curated in R Core (R version 4.1.1), (R Core Team, 2021). The iAUC was integrated using the *mosaicCalc* (version 0.5.1), (Kaplan D.T., Pruim R., & Horton N.J., 2020) package. Annotated code and source data are available as Supplementary.

## Results

Dough formulation was similar for *sbeII* and WT control breads, however *sbeII* flour required additional water (39% for *sbeII* and 37% for WT control) to produce a workable dough. After mixing, the dough was assessed visually and found to be well developed. Both roll types developed and baked well, reaching a core temperature of 95°C.

Rolls were made to deliver an equal starch content; the *sbeII* flour was characterised by a lower total starch content, therefore the *sbeII* rolls were ~10 g larger than the WT control rolls (161.26 g ± 0.16 g, 155.73 g ± 0.40 g, respectively, mean ± SEM, n=3).

### Moisture

Moisture changes are shown in Figure 2, data is shown in Supplementary Table 2. During baking, moisture loss of bread rolls made with *sbeII* flour was 11.9% compared to 10.3% in WT controls.

**Figure 2.**
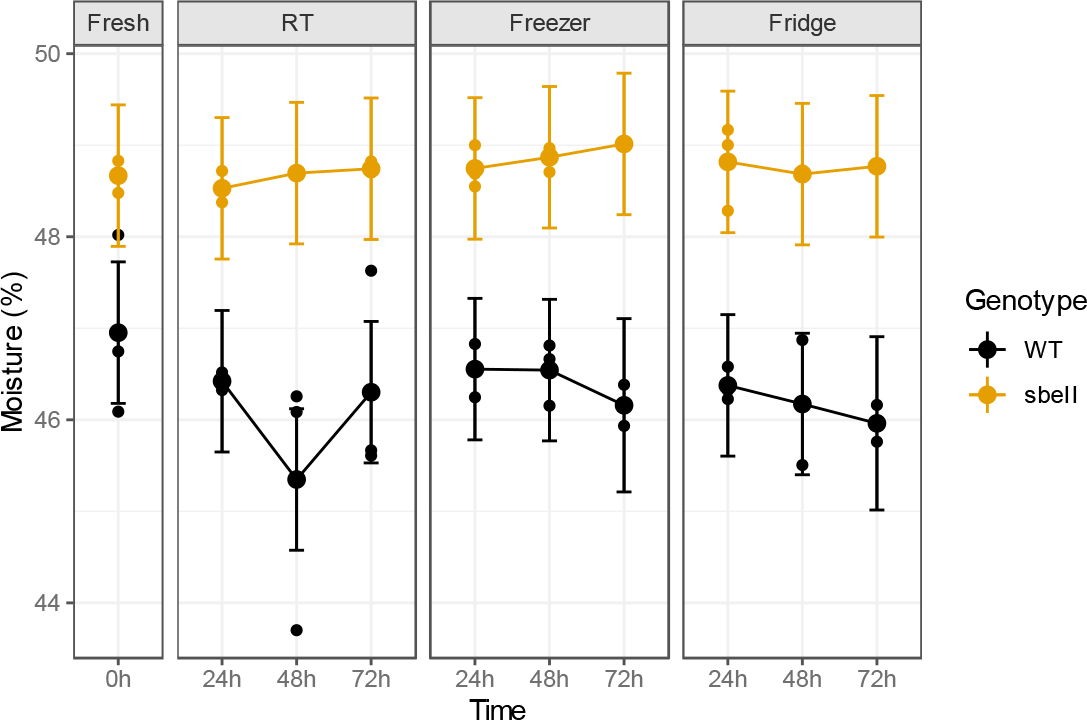
Estimated marginal means of moisture during storage, n = 3 with error bars representing the 95%CI.

Freshly baked *sbeII* bread rolls had a small but higher moisture content (48.7%) compared to WT breads (47%; *p* = 0.003). The moisture difference between genotypes was maintained consistently over storage, regardless of the storage conditions (interaction between genotype and storage *p* = 0.3, 0.1, 0.1 for RT, Freezer and Fridge).

### Starch digestibility in bread

Starch digestibility was defined using the following parameters: the extent of digestibility within 90 min of incubation (*C_90_*), the predicted starch digested at the end of reaction (C_*∞*_), the digestibility rate constant (*k*) and incremental area under the curve (iAUC), Figure 3D-G.

**Figure 3.**
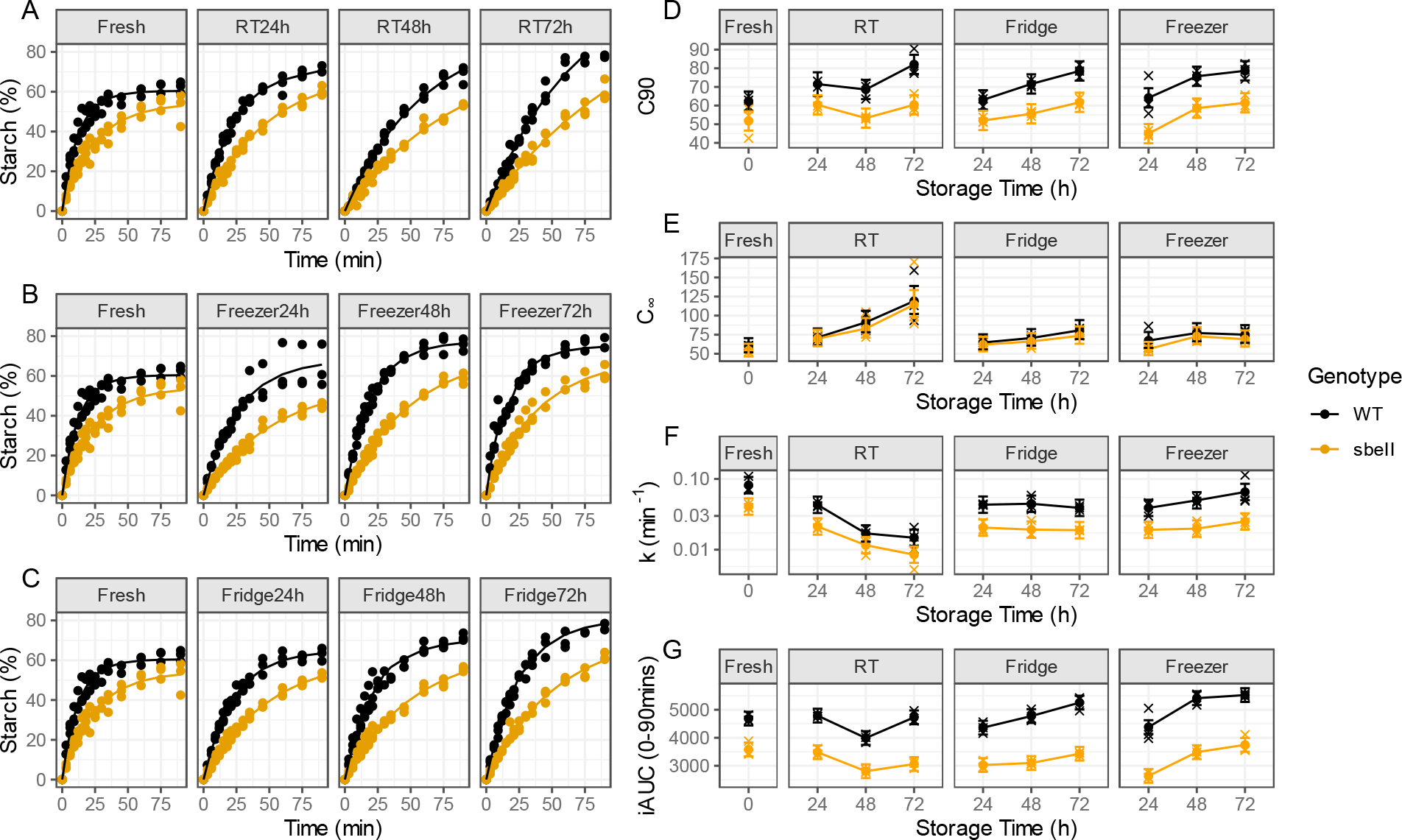
**A-C.** Starch digestibility curves of sbeII (yellow) and WT control (black) bread samples, experimental data points represent independently treated samples, n = 3 per condition. Experimental data (replicate datapoints) are shown by fitting a first-order equation based on the estimates of k and C_∞_ values (n = 3 independent samples) obtained from a non-linear regression model. **D-G**. Grouped means of parameters obtained from digestibility curves, (D) C_90_, (E) C_∞_, (F) k, (G) iAUC, error bars represent the 95%CI (n = 3).

Overall, starch in *sbeII* fresh bread was significantly less digested than the WT control, as shown by the *C_90_* and iAUC (*p* = 0.005, *p* = <0.001, respectively), regardless of storage, Supplementary Table 3.

Compared to freshly baked bread, storage led to increased starch digested at 90 minutes (C_*90*_, *p* = <0.001, for all conditions), in both bread genotypes, but there was no evidence of an interaction effect between storage and genotype on starch digested (C_*90*_, *p* = 0.3), showing that the effect difference between *sbeII* and WT control bread is maintained throughout storage.

The extent of starch digestibility (C_*90*_) was 51.7% in *sbeII* freshly baked bread compared to 61.4%, 61.7% and 60.4% when *sbeII* bread was stored for 72h in the freezer, fridge and RT respectively. In freshly baked WT control bread, starch digestibility was 62.4% and increased to 78.6%, 78.5% and 82.1% after 72h freezer, fridge and RT storage, respectively. For samples stored at RT, some evidence of interaction between storage and bread genotype was observed (p = 0.03). Here, C_*90*_ was higher after 48h storage at RT compared to fresh bread.

These changes are relative to the starch digested within 90 minutes of incubation however, the model suggests that digestion of *sbell* bread is slower compared to the WT control, and at the end of the reaction the predicted differences between the breads would be negligible (as evidenced by estimates of the C_*∞*_ parameter), (Figure 3E).

The digestion rate (*k*) of starch in *sbeII* bread was overall lower compared to starch in WT control bread (*p* = <0.001) and no effect of interaction between genotype and storage was found (p = 0.1). Compared to fresh bread, RT and fridge storage affected the rate of starch digestion in both bread (*p* = <0.001, both conditions) but not in breads stored in the freezer (*p* = 0.2), Figure 3A-C.

Considering the iAUC of digestibility curves, besides the significant effect of the genotype (*p* = <0.001), there was a significant effect of the fridge and freezer storage on iAUC, compared to fresh bread (*p* = 0.002 and *p* = <0.001, respectively). There was some evidence of an interaction effect between storage and genotype on iAUC however, the magnitude of the effect is relatively small as the iAUC summarises rate (*k*) and extent of digestibility (C_*90*_), (*p* = 0.0007).

The estimated marginal means with 95% CI for all storage conditions are reported in Supplementary Table 3.

### Texture analysis

Macrostructure of bread crumb was analysed instrumentally for textural differences between bread types, during storage. Estimated means of all characteristics following each storage conditions are reported in Supplementary Table 4 and are showed in Figure 4.A.

**Figure 4.**
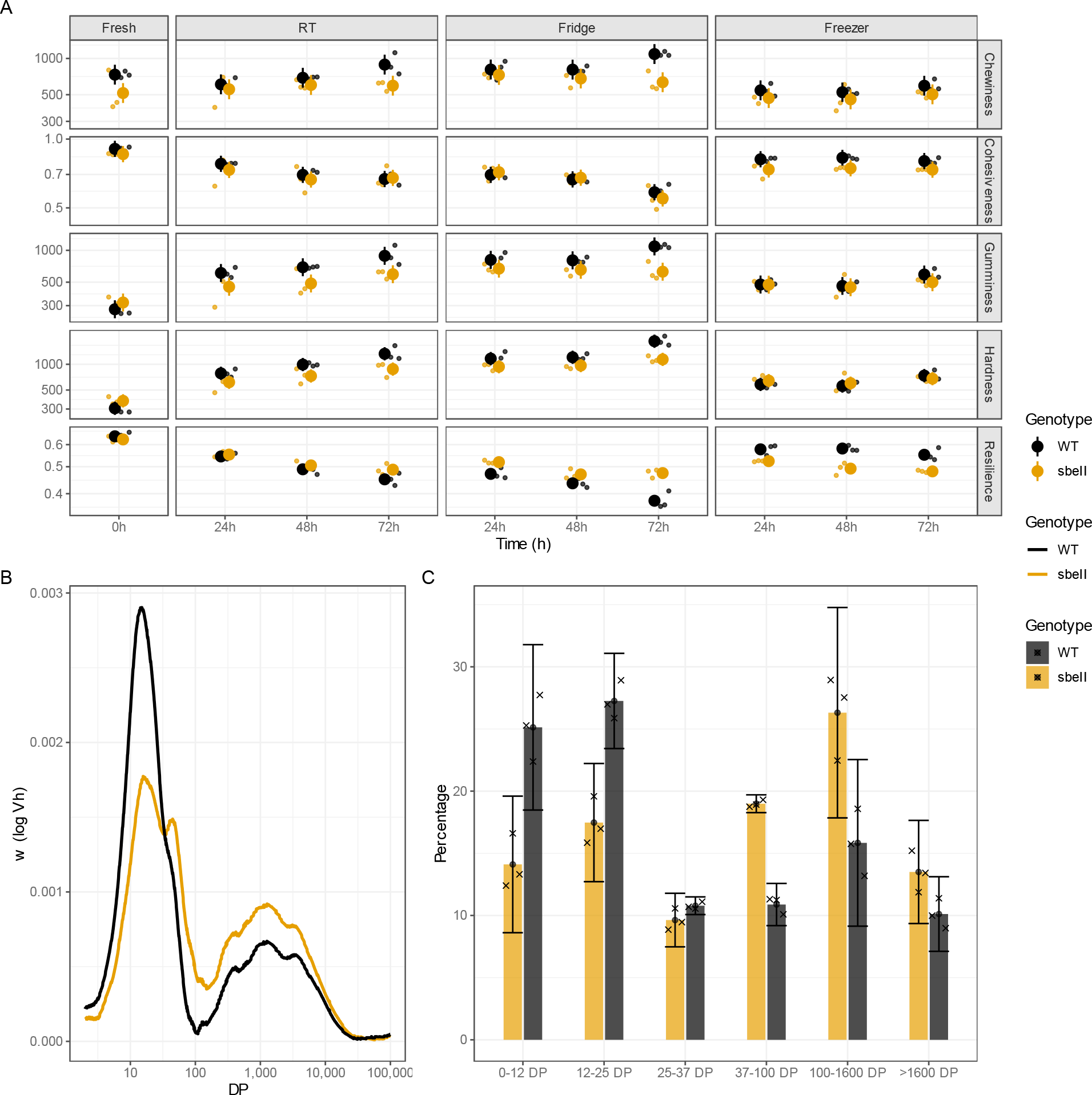
A. Texture profile of WT and sbeII breads, data points represent the mean value with error bars = ± 95% CI, n = 3 independent bread rolls were analysed per condition, with four technical replicates per bread roll. **B**. Chain-length distribution of debranched sbeII and WT control starch isolated from flour determined by SEC, average of n = 3 independent replicates. **C**, Fractions grouped, with error bars = 95% CI, based on DP as described by Hanashiro et al.(Hanashiro, et al., 1996) and Vilaplana et al. (Vilaplana, et al., 2012) where amylopectin short chains are 0-37 DP, amylopectin long chains are 37-100 DP, amylose short chains are 100-1600 DP and amylose long chains are >1600 DP.

Freshly baked *sbeII* bread was similar to the WT control for hardness (*sbeII* = 373 g and WT = 306 g, *p* = 0.1), cohesiveness (*sbeII* = 0.859%, and WT = 0.906 g, *p* = 0.3), resilience (*sbeII* = 0.62% and WT = 0.64%, *p* = 0.4) but showed lower chewiness compared to the WT control (*sbeII* = 516 and WT = 734, *p* = 0.01).

With storage, increased crumb firming and decreased resilience was observed in both bread genotypes. Firming was less pronounced in *sbeII* breads than in the WT control while loss of resilience was more marked in *sbeII* bread than the WT control (*p* = <0.001, both parameters). Overall, the effect of storage on crumb hardness and resilience varied by genotype (storage by genotype interaction effect on hardness *p* = 0.001, effect on resilience *p* = <0.001) but not when considering bread chewiness and cohesiveness.

There was less effect on bread hardness with freezing, with no difference in the effect of freezing between genotypes.

Following storage, the *sbell* bread appeared slightly chewier than the WT, with the most marked difference after 72h in the fridge or at RT (*sbeII* fridge 72h = 634.5, WT fridge 72h = 1088, *p* = 0.0002, *sbeII* RT 72h = 593.9, WT RT 72h = 888.9, *p* = 0.004).

Resilience of *sbeII* breads was marginally lower than the WT control after 24h of freezer storage (*sbeII* freezer 24h = 0.52, WT freezer 24h = 0.57, *p* = 0.003) but slightly higher than the WT control after fridge storage (*sbeII* fridge 24h = 0.51% and WT fridge 24h = 0.47, *p* = 0.004).

### Chain length distribution

Analysis of starch chain-length distribution showed differences in the molecular structure of amylose and amylopectin from *sbeII* mutant wheat compared to the WT control.

The proportion of long amylopectin chains (DP 37 to 100) was higher in *sbeII* starch (19% ± 0.3%) compared to the WT control (10.9% ± 0.7%) and presented two distinct peaks (Figure 4, B), in contrast to a slight shoulder peak in WT starch. The *sbeII* starch was also characterised by a larger proportion of amylose chains (DP 100 – 1600) compared to the WT control (26.3% ± 3.4, 15.8% ± 2.7%, respectively). The percentage of short chains (DP<25) in mutants was lower in the *sbeII* mutant compared to the WT control (Figure 4, C).

Chain length proportions per DP fraction are reported as Mean ± SD, n = 3. All fractions can be found in Supplementary Table 5.

### Microscopy

Micrographs of starch in water using regular brightfield microscopy complemented by polarised-light microscopy were used to indirectly assess starch crystallinity in raw flour and in bread.

Starch in raw *sbeII* flour showed a typical birefringence pattern (‘Maltese cross’) however, this was less defined in starch granules with altered morphology Figure 5.A.

**Figure 5.**
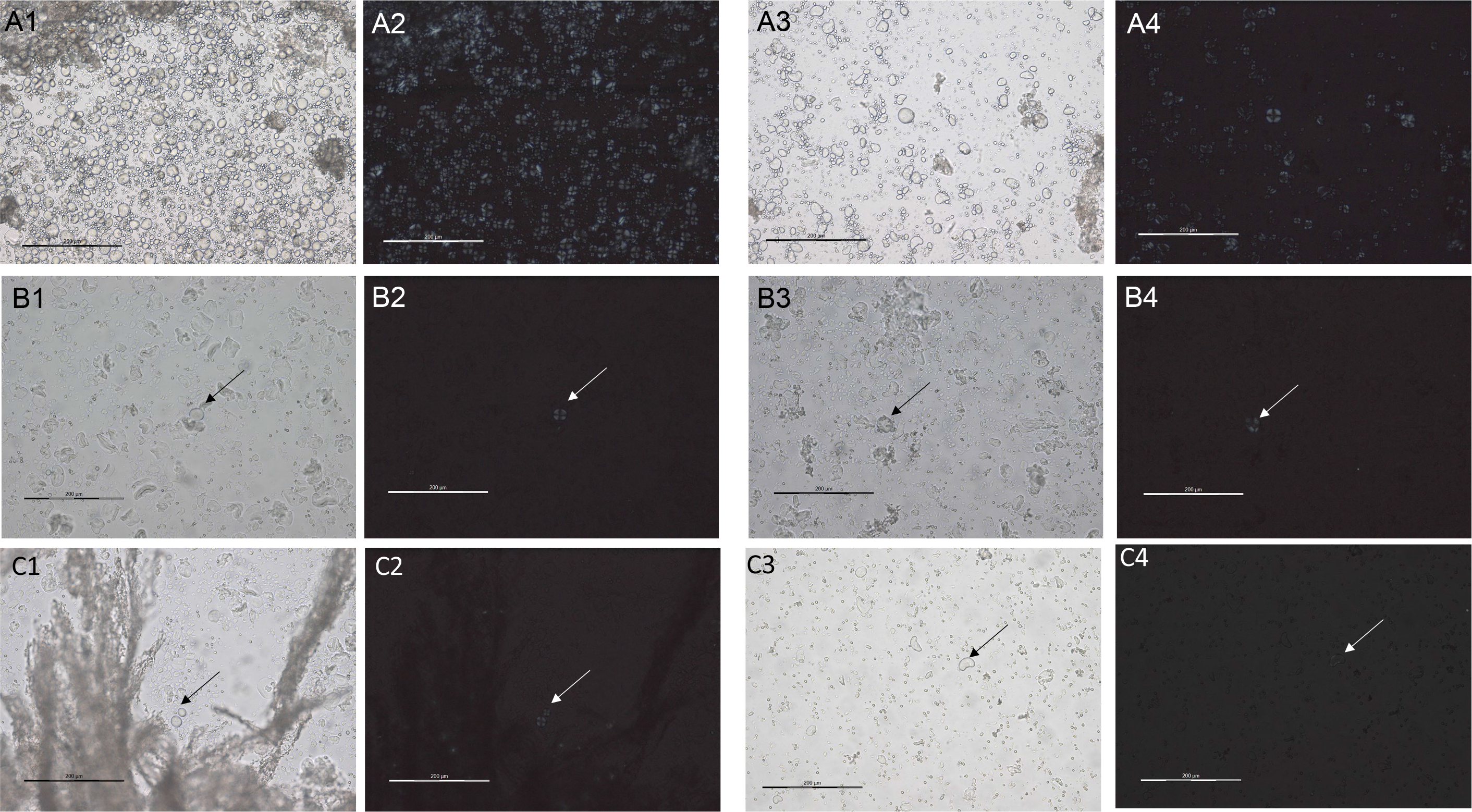
Micrographs of starch in water using brightfield (BF) and polarised light filter (P) highlighting birefringence of starch granules. **1**. WT control starch under BF; **2**. WT control starch under P; **3**. sbeII mutant starch under BF; **4**. sbeII mutant starch under P; **A**. starch from flour; **B**. Starch from fresh bread crumb; **C**. Starch from bread crumb after 7 days of freezer storage. Scale bar in white or black represent 200 μm.

In freshly baked bread, only few starch granules in both WT and *sbeII* crumb were visibly birefringent suggesting loss of starch semi-crystalline structure during baking (Figure 5.B).

Breads were then stored for 7 days in the freezer and thawed once. A larger number of granules with a birefringent pattern was observed in the crumb WT control bread, while almost no granules with such pattern were visible in the crumb of frozen *sbeII* bread, Figure 5.C.

The crust of breads was also included in the analysis. Here, the birefringence was less evident in *sbeII* starch granules from fresh crust compared to the WT control granules, possibly due to different levels of gelatinisation and/or crystallinity (Supplementary Figure 2.D). Granules with birefringent pattern were also observed in the crust of frozen bread, both mutant *sbeII* and WT control, however to a lower extent (Supplementary Figure 2.E).

## Discussion

The use of flour with higher-amylose content than conventional wheat in the production of starch-based foods has the potential to elicit a lower glycaemic response compared to conventional starch-based foods (Ang K., et al., 2020; Granfeldt Y., Drews A., & Björck I., 1995; Hallström E., et al., 2011; Hoebler C., Karinthi A., Chiron H., Champ M., & Barry J.L., 1999). In this study we investigated the macro- and microstructural changes in bread made from a high-amylose wheat flour during three days of storage. The *sbeII* starch in flour had a higher proportion of intermediate chains (DP 37-100) compared to the WT control, as a result of the reduced activity of SBEII enzymes during starch biosynthesis. This is consistent with results from other studies of crops with mutations in *starch branching enzyme* genes (Jane J., et al., 1999; Regina A., et al., 2015).

Starch in white bread made from *sbeII* mutant wheat flour was found to be less susceptible to amylolysis when fresh and after storage, and it was characterised by a consistently slower digestion rate (*k*) leading to lower starch digested (iAUC). Therefore, the lower starch digestibility observed in *sbeII* fresh and frozen breads may be due to enzyme-resistant structures already present after baking and linked to the amylose content rather than structures formed during storage.

Considering the different digestibility parameters, we have shown that the percentage of starch digested in *sbeII* bread at 90 mins is consistently lower than the starch digestibility from WT control bread, but the rate starch hydrolysis can be affected by the storage condition.

Overall, the effect of the storage condition can affect *sbeII* starch digestibility early on during storage compared to the WT control, however the difference between *sbeII* bread and WT control is maintained during storage. This could be due to the increased amylose proportion in *sbeII* starch promoting faster retrogradation compared to the WT control. This was also observed in microscopy images, where *sbeII* starch granules in fresh bread and after 7 days of storage in freezer, did not show the birefringence pattern typical of recrystallised granules, as in the WT control.

A recent study measured RS content of bread during storage and reported that bread made from partial substitution (50%) of conventional white flour with high-amylose wheat flour reaching 51.5% amylose content, and stored in the fridge for up to 5 days (Li C. & Gidley M.J., 2022). Interestingly, they did not observe an increase in RS content of bread during storage. We observed the starch in *sbeII* bread remained less susceptible to amylolysis compared to the WT control. When left at RT, the rate of starch hydrolysis of both bread types decreased within three days of baking, compared to fresh bread while that starch digestibility (C_*90*_ and iAUC) in bread stored in the fridge increased over time compared to fresh bread however, this may be due to variation in the moisture content of samples stored. The study design did not allow to determine moisture on ground bread crumb samples at each storage timepoint. This was due to time constraints in carrying out the digestibility experiments over several days with the different storage conditions and a sufficient number of replicates. Therefore, moisture was measured on fresh samples only (when bread sampling). This could have led to overestimation of starch digestibility for the samples stored and it is a limitation of the study.

However, since both *sbeII* and WT control breads have been treated the same way, the effect size measured in this study is not affected. The single enzyme assay used to determine starch digestibility here is a sensitive method and may be able to detect subtle differences compared to the RS AOAC method by producing several experimental parameters to characterise the digestibility curves providing accurate estimates of digestion. The percentage of starch hydrolysed by α-amylase, as reported in this study (C_*90*_), has been shown to correlate well with glycaemic index measurements in human participants (Edwards C.H., et al., 2019), suggesting that the lower starch digestibility of *sbeII* starch in bread has potential for applications in frozen bread, as shown by a previous study (Corrado M., et al., 2022), but also when bread is consumed fresh, or after storage in the fridge or at room temperature.

The higher hardness of *sbeII* fresh bread compared to the WT control is has been attributed to the presence of increased amylose content in *sbeII* starch (Goesaert, Slade, Levine, & Delcour, 2009) and its ability to form double helices or aggregates of helices. Research in gluten free breads using high amylose rice flour has shown that lower amylose content rice types led to softer crumb (Roman, Reguilon, Gomez, & Martinez, 2020), like the softer freshly baked WT control bread in this study when compared to the freshly baked *sbeII* bread. Future research studies investigating structural organisation of starch chains may help clarify this.

Based on the texture measurements reported in this study, macrostructural changes to the crumb texture set in earlier during storage in *sbeII* bread compared to the WT control. Overall, texture changes in *sbeII* bread crumb were limited compared to the WT control crumb, suggesting a more stable structure over storage. This may be due partly to the water, as a crystallized rigid starch network formed during staling can trap bound water favouring crumb firmness (Hemdane, et al., 2017). While moisture of bread as a whole did not vary significantly during storage, migration of water from the protein to the starch fraction may have led to the strengthening of the starch network, favouring water retention and greater crumb firmness (Bosmans G. M., Lagrain B., Ooms N., Fierens E., & Delcour J. A., 2013), though water migration was not examined in detail in this study. However, the marginally higher water content of *sbeII* breads, due to the known higher water holding capacity of the *sbeII* flour, may have contributed to the limited crumb firming within the first three days of storage. As previously shown, moisture levels between 40%-50%, as the breads in this study, are ideal for starch crystallisation as it promotes network plasticisation and favouring retrogradation (Zeleznak & Hoseney, 1986). Similarly, Li and Gidley (Li C. & Gidley M.J., 2022) showed that substitution of conventional wheat white flour with 50% high-amylose (51.5%) flour in bread making does not lead to a noticeable increase in bread firmness over storage in the fridge. Barros *et al*., reported that 10%-20% wheat flour replacement with HI-MAIZE 260® in bread making delayed bread staling by decreasing the rate of starch recrystallisation, without a negative impact on bread quality (Barros J. H. T., Telis V. R. N., Taboga S., & Franco C. M. L., 2018). Evident changes in crumb firmness have been shown to be linked to amylopectin retrogradation, especially at freezer temperatures, (Bosmans G. M., Lagrain B., Fierens E., et al., 2013) and independent from the moisture loss (Ribotta P.D., et al., 2003).

These studies support the findings reported here; considering the limited micro- and macrostructural changes in bread made from *sbeII* flour during storage, this high-amylose flour could be used to make bread with lower glycaemic potency for commercial use thanks to the limited staling.

Interestingly, instrumentally measured resilience (a measure of elasticity) of *sbeII* bread crumb was found to be slightly higher compared to the WT control after storage in the fridge but a little lower than the control after storage in the freezer, suggesting higher stability of refrigerated *sbeII* bread compared to conventional bread.

This is promising for potential use of this wheat in production of sandwich bread as it is common to use dough improvers to decrease the progression of staling and reduce bread firming (Hug-Iten, Escher, & Conde-Petit, 2003). In this study, the formulation, particularly the amount of water used to hydrate the flour and form the dough, may have played a role in promoting bread softness over time. In a recent publication (Arp C.G., et al., 2020) it was reported that in breads with intermediate content of resistant starch (10% and 20% high-amylose flour replacement), the good initial quality and slower dehydration rate of the matrix counteracted the negative effects of starch retrogradation on bread texture. While no major water adjustments were made to form the *sbeII* dough, the WT control dough was slightly stickier than the *sbeII* dough. The dough produced with both flour types were well developed. It is noteworthy that the bread making method used here was closer to home baking than the industrial Chorleywood method, so bread texture may differ when scaling up the formulation for large industrial production of convenience bread. Based on results from this study, the *sbeII* wheat flour shows promise for home and artisan bread making with lower starch digestibility, however other studies on dough development and machinability are required.

## Conclusions

Bread made from *sbeII* wheat flour was found to be more stable during storage and less prone to staling than the WT control, representing conventional white bread. This is likely due to the increase in amylose proportion in *sbeII* starch. Despite requiring more water to form a well-developed dough compared to conventional wheat white flour, as shown by previous research, the starch digestibility of *sbeII* bread remained consistently lower than the WT control over storage. At the texture level, *sbeII* bread was less prone to firming compared to WT control bread.

This shows potential for *sbeII* bread to be used to develop freshly prepared and refrigerated wheat bread with a lower glycaemic potency and similar or, in some cases, better ageing than conventional white bread.

## Authors contribution

Marina Corrado: Conceptualization, investigation, methodology, project administration, data curation and formal analysis, writing – original draft preparation, writing – review & editing, Petros Zafeiriou: investigation, methodology, writing – review & editing, Jennifer Ahn-Jarvis: investigation, methodology, writing – review & editing, George M. Savva: methodology, data curation, visualisation, formal analysis, writing – review & editing, Cathrina H. Edwards: conceptualization, methodology, supervision, writing – review & editing, Brittany A. Hazard: conceptualization, funding acquisition, methodology, supervision, writing – review & editing.

