## Supplementary Information for "Impact of storage on starch digestibility and texture of a high-amylose wheat bread"

*Table 1 Estimated nutritional content of breads and indirect damaged starch measured by Solvent Retention Capacity assay.*

| **Proximate measures** | **WT bread** | ***sbeII* bread** |
|---|---|---|
| Moisture (g/100g) | 46 | 48 |
| Protein (g/100g) | 8.8 | 8.2 |
| Total fat (g/100g) | 3.7 | 2.5 |
| Saturated fat (g/100g) | 1.1 | 0.8 |
| Monounsaturated fat (g/100g) | 0.2 | 0.1 |
| Polyunsaturated fat (g/100g) | 0.7 | 0.5 |
| Total Carbohydrate (g/100g) | 37.8 | 37.7 |
| Starch (g/100g) | 41.0 | 39.4 |
| Total sugars (g/100g) | 11.0 | 12.0 |
| Sucrose (g/100g) | 0.4 | 0.3 |
| Ash (g/100g) | 0.7 | 0.7 |
| AOAC Fibre (g/100g) | 2.9 | 1.8 |
| Energy (kcal/100g) | 214.1 | 203.8 |
| Energy (kJ/100g) | 906.8 | 863.4 |
| Sodium (mg/100g) | 1022.5 | 1030.0 |
| Potassium (mg/100g) | 95.0 | 93.1 |
| Calcium (mg/100g) | 17.5 | 15.0 |
| Magnesium (mg/100g) | 26.6 | 17.4 |
| Phosphorus (mg/100g) | 105.5 | 80.4 |
| Iron (mg/100g) | 0.7 | 0.5 |
| Zinc (mg/100g) | 0.7 | 0.6 |
| Manganese (mg/100g) | 0.6 | 0.5 |
| Indirect damaged starch* ($Na_2CO_3$ absorption) | 74.8 | 86.8 |

**Briefly, 5g of flour are suspended in 25 mL of 5% (w/w or v/v) of Na2CO3. Samples are left to hydrate for 20 minutes with intermittent mixing, followed by centrifugation at 1000 x g for 15 minutes. The supernatant is discarded and the tubes are left 10 minutes to dry upside down on paper. The pellet weight is recorded to calculate solvent retention capacity (SRC) as follows:*

*SRC value (%)= [[(gel wt/flour wt) × (86/(100 − % flour moisture)) − 1)] × 100] (Kweon, Slade, & Levine, 2011).*

*Table 2. Weight of bread rolls before and after baking, and after storage. Moisture measured by air-oven method. Mean ± SEM, n=3 independent samples per condition.*

| Time (h) | Storage | Genotype | Moisture (%) | Dough weight (g) | Baked roll weight (g) | Roll weight after storage (g) |
|---|---|---|---|---|---|---|
| 0 | Fresh | *sbeII* | 48.76 ± 0.12 | 183.06 ± 0.21 | 161.26 ± 0.16 | - |
| 0 | Fresh | WT | 46.50 ± 0.20 | 173.56 ± 0.36 | 155.73 ± 0.40 | - |
| 24 | Freezer | *sbeII* | 48.74± 0.13 | 183.13 ± 0.14 | 160.20 ± 0.39 | 160.13 ± 0.41 |
| 24 | Freezer | WT | 46.55 ± 0.16 | 173.73 ± 0.28 | 156.03 ± 0.49 | 156.03 ± 0.49 |
| 24 | Fridge | *sbeII* | 48.81 ± 0.27 | 183.03 ± 0.18 | 160.66 ± 0.31 | 160.63 ± 0.29 |
| 24 | Fridge | WT | 46.37 ± 0.10 | 173.60 ± 0.30 | 155.06 ± 0.66 | 155.06 ± 0.63 |
| 24 | RT | *sbeII* | 48.52 ± 0.10 | 183.13 ± 0.18 | 161.40 ± 0.20 | 161.30 ± 0.29 |
| 24 | RT | WT | 46.42 ± 0.05 | 173.70 ± 0.20 | 156.06 ± 0.12 | 155.90 ± 0.29 |
| 48 | Freezer | *sbeII* | 48.86 ± 0.08 | 183.20 ± 0.11 | 161.90 ± 0.30 | 161.86 ± 0.21 |
| 48 | Freezer | WT | 46.54± 0.19 | 173.33 ± 0.24 | 154.53 ± 0.23 | 154.53 ± 0.23 |
| 48 | Fridge | *sbeII* | 48.68 ± 0.02 | 183.20 ± 0.15 | 161.23 ± 0.64 | 161.23 ± 0.58 |
| 48 | Fridge | WT | 46.17 ± 0.39 | 173.50 ± 0.28 | 154.73 ± 0.46 | 154.73 ± 0.41 |
| 48 | RT | *sbeII* | 48.69 ± 0.01 | 183.16 ± 0.14 | 161.63 ± 0.43 | 161.33 ± 0.48 |
| 48 | RT | WT | 45.34 ± 0.82 | 173.30 ± 0.20 | 155.26 ± 0.46 | 155.03 ± 0.43 |
| 72 | Freezer | *sbeII* | 49.01 ± 0.00 | 183.03 ± 0.31 | 160.80 ± 0.55 | 160.70 ± 0.55 |
| 72 | Freezer | WT | 47.49 ± 1.34 | 174.16 ± 0.29 | 155.70 ± 0.70 | 155.70 ± 0.70 |
| 72 | Fridge | *sbeII* | 48.76 ± 0.01 | 183.03 ± 0.26 | 160.83 ± 0.48 | 160.83 ± 0.48 |
| 72 | Fridge | WT | 47.58 ± 1.62 | 174.36 ± 0.31 | 156.13 ± 0.64 | 156.10 ± 0.64 |
| 72 | RT | *sbeII* | 48.74 ± 0.04 | 182.96 ± 0.29 | 161.63 ± 0.35 | 161.36 ± 0.32 |
| 72 | RT | WT | 46.30 ± 0.66 | 174.10 ± 0.47 | 156.10 ± 0.92 | 155.80 ± 0.92 |

*Table 3. Starch susceptibility to amylolysis in bread. Estimated marginal means with 95% CI, n = 3 independent samples per condition. Endogenous maltose present before starting the reaction (Y0), first order rate constant (k) and predicted starch digested at the end of reaction (C∞), estimated after subtracting Y0 from the subsequent timepoints. $C_{90}$ (experimental endpoint) and the Area Under the Curve (AUC)*

| Genotype | Storage | Time | Y0 (%) | $C_{90}$ (%) | AUC (%min$^{-1}$) | $k$ (min$^{-1}$) | $C_\infty$ (%) |
|---|---|---|---|---|---|---|---|
| *sbeII* | Fresh | 0h | 14.8 [13.5; 16.1] | 51.7 [46.6; 56.9] | 3581 [3336; 3826] | 0.040 [0.028; 0.053] | 54.2 [37.6; 70.8] |
| | RT | 24h | 12.1 [10.8;13.3] | 60.3 [55.2; 65.5] | 3482 [3237; 3727] | 0.021 [0.008; 0.033] | 69.9 [53.3; 86.5] |
| | RT | 48h | 14.1 [12.7; 15.4] | 53.3 [48.1; 58.4] | 2805 [2560; 3050] | 0.011 [-0.0004; 0.024] | 84.1 [67.6; 100.7] |
| | RT | 72h | 11.8 [10.5; 13.1] | 60.4 [55.2; 65.6] | 3060 [2815; 3305] | 0.009 [-0.003; 0.021] | 119.4 [102.8; 135.9] |
| | Freezer | 24h | 9.8 [8.4; 11.1] | 44.9 [39.8; 50.1] | 2636 [2391; 2881] | 0.019 [0.006; 0.031] | 56.1 [39.6; 72.7] |
| | Freezer | 48h | 12.3 [10.9; 13.6] | 58.6 [53.4; 63.8] | 3486 [3240; 3731] | 0.020 [0.007; 0.032] | 73.1 [56.5; 89.7] |
| | Freezer | 72h | 12.6 [11.2; 13.9] | 61.4 [56.3; 66.6] | 3746 [3501; 3991] | 0.025 [0.012; 0.037] | 69.1 [52.5; 85.6] |
| | Fridge | 24h | 10.7 [9.4; 12.1] | 52.0 [46.8; 57.1] | 3025 [2780; 3271] | 0.020 [0.008; 0.033] | 62.0 [45.4; 78.5] |
| | Fridge | 48h | 13.7 [12.4; 15.1] | 55.6 [50.4; 60.7] | 3097 [2852; 3343] | 0.019 [0.007; 0.032] | 66.5 [49.9; 83.1] |
| | Fridge | 72h | 12.4 [11.1; 13.8] | 61.7 [56.6; 66.9] | 3429 [3184; 3674] | 0.018 [0.006; 0.031] | 73.5 [57.0; 90.1] |
| WT | Fresh | 0h | 11.7 [10.4; 13.0] | 62.4 [57.2; 67.6] | 4682 [4437; 4927] | 0.083 [0.071; 0.095] | 60.4 [43.9; 77.0] |
| | RT | 24h | 11.6 [10.2; 12.9] | 71.5 [65.1; 77.9] | 4789 [4544; 5034] | 0.043 [0.030; 0.055] | 71.3 [54.7; 87.8] |
| | RT | 48h | 12.5 [11.2; 13.9] | 68.6 [63.5; 73.8] | 3989 [3744; 4234] | 0.017 [0.004; 0.029] | 91.6 [75.0; 108.2] |
| | RT | 72h | 11.1 [9.7; 12.4] | 82.1 [76.9; 87.2] | 4727 [4482; 4972] | 0.015 [0.002; 0.027] | 122.0 [105.5; 138.6] |
| | Freezer | 24h | 9.4 [8.0; 10.7] | 64.1 [59.0; 69.3] | 4380 [4134; 4625] | 0.039 [0.027; 0.052] | 68.4 [51.8; 84.9] |
| | Freezer | 48h | 12.7 [11.4; 14.1] | 75.7 [70.5; 80.8] | 5412 [5167; 5657] | 0.049 [0.037; 0.062] | 77.2 [60.7; 93.8] |
| | Freezer | 72h | 14.2 [12.8; 15.5] | 78.6 [73.5; 83.8] | 5521 [5276; 5767] | 0.070 [0.058; 0.083] | 75.2 [58.7; 91.8] |
| | Fridge | 24h | 10.1 [8.8; 11.5] | 63.0 [57.9; 68.2] | 4359 [4114; 4605] | 0.043 [0.030; 0.055] | 64.7 [48.2; 81.3] |
| | Fridge | 48h | 13.0 [11.7; 14.3] | 71.6 [66.4; 76.7] | 4773 [4527; 5018] | 0.045 [0.033; 0.057] | 70.6 [54.0; 87.2] |
| | Fridge | 72h | 12.4 [11.1; 13.8] | 78.5 [73.4; 83.7] | 5253 [5008; 5498] | 0.039 [0.026; 0.051] | 80.7 [64.1; 97.3] |

*Table 4. Texture analysis measurements, Estimated marginal means with 95% CI, n = 3 independent samples per condition (4 technical replicates)*

| Storage Condition | Storage duration | Genotype | Response | SE | df | lower.CL | upper.CL | Texture measure |
|---|---|---|---|---|---|---|---|---|
| Fresh | 0h | WT | 305.8183 | 26.939 | 40 | 223.961 | 417.5942 | Hardness |
| Fresh | 0h | *sbeII* | 373.4318 | 32.895 | 40 | 273.4767 | 509.9203 | Hardness |
| Freezer | 24h | WT | 581.705 | 51.241 | 40 | 426.0021 | 794.3169 | Hardness |
| Freezer | 24h | *sbeII* | 642.7244 | 56.616 | 40 | 470.6887 | 877.6388 | Hardness |
| Fridge | 24h | WT | 1164.528 | 102.58 | 40 | 852.8233 | 1590.161 | Hardness |
| Fridge | 24h | *sbeII* | 936.6819 | 82.51 | 40 | 685.9637 | 1279.037 | Hardness |
| RT | 24h | WT | 783.4469 | 69.012 | 40 | 573.7445 | 1069.795 | Hardness |
| RT | 24h | *sbeII* | 616.8406 | 54.336 | 40 | 451.7332 | 842.2945 | Hardness |
| Freezer | 48h | WT | 556.6381 | 48.742 | 39 | 408.1547 | 759.1387 | Hardness |
| Freezer | 48h | *sbeII* | 600.6628 | 52.911 | 40 | 439.8856 | 820.2038 | Hardness |
| Fridge | 48h | WT | 1207.639 | 106.38 | 40 | 884.3946 | 1649.028 | Hardness |
| Fridge | 48h | *sbeII* | 965.9951 | 85.092 | 40 | 707.4307 | 1319.064 | Hardness |
| RT | 48h | WT | 992.0864 | 87.391 | 40 | 726.5382 | 1354.692 | Hardness |
| RT | 48h | *sbeII* | 729.762 | 64.283 | 40 | 534.4292 | 996.4884 | Hardness |
| Freezer | 72h | WT | 736.7416 | 64.898 | 40 | 539.5406 | 1006.019 | Hardness |
| Freezer | 72h | *sbeII* | 679.0834 | 59.819 | 40 | 497.3156 | 927.2869 | Hardness |
| Fridge | 72h | WT | 1860.303 | 163.87 | 40 | 1362.362 | 2540.239 | Hardness |
| Fridge | 72h | *sbeII* | 1141.099 | 100.52 | 40 | 835.665 | 1558.168 | Hardness |
| RT | 72h | WT | 1326.348 | 116.84 | 40 | 971.329 | 1811.125 | Hardness |
| RT | 72h | *sbeII* | 878.3376 | 77.371 | 40 | 643.2362 | 1199.368 | Hardness |
| Fresh | 0h | WT | 0.906002 | 0.0365 | 40 | 0.785811 | 1.044576 | Cohesiveness |
| Fresh | 0h | *sbeII* | 0.859134 | 0.0346 | 40 | 0.74516 | 0.99054 | Cohesiveness |
| Freezer | 24h | WT | 0.816017 | 0.0329 | 40 | 0.707763 | 0.940828 | Cohesiveness |
| Freezer | 24h | *sbeII* | 0.736312 | 0.0296 | 40 | 0.638632 | 0.848932 | Cohesiveness |
| Fridge | 24h | WT | 0.697355 | 0.0281 | 40 | 0.604843 | 0.804016 | Cohesiveness |
| Fridge | 24h | *sbeII* | 0.7163 | 0.0288 | 40 | 0.621275 | 0.825859 | Cohesiveness |
| RT | 24h | WT | 0.779263 | 0.0314 | 40 | 0.675885 | 0.898453 | Cohesiveness |
| RT | 24h | *sbeII* | 0.732887 | 0.0295 | 40 | 0.635662 | 0.844984 | Cohesiveness |
| Freezer | 48h | WT | 0.82841 | 0.0324 | 35 | 0.720284 | 0.952768 | Cohesiveness |

| | | | | | | | | |
|---|---|---|---|---|---|---|---|---|
| Freezer | 48h | *sbeII* | 0.745085 | 0.03 | 40 | 0.646242 | 0.859048 | Cohesiveness |
| Fridge | 48h | WT | 0.665942 | 0.0268 | 40 | 0.577598 | 0.7678 | Cohesiveness |
| Fridge | 48h | *sbeII* | 0.677572 | 0.0273 | 40 | 0.587684 | 0.781208 | Cohesiveness |
| RT | 48h | WT | 0.697285 | 0.0281 | 40 | 0.604783 | 0.803937 | Cohesiveness |
| RT | 48h | *sbeII* | 0.665358 | 0.0268 | 40 | 0.577091 | 0.767126 | Cohesiveness |
| Freezer | 72h | WT | 0.801086 | 0.0323 | 40 | 0.694813 | 0.923614 | Cohesiveness |
| Freezer | 72h | *sbeII* | 0.734578 | 0.0296 | 40 | 0.637128 | 0.846933 | Cohesiveness |
| Fridge | 72h | WT | 0.585278 | 0.0236 | 40 | 0.507635 | 0.674798 | Cohesiveness |
| Fridge | 72h | *sbeII* | 0.549974 | 0.0221 | 40 | 0.477014 | 0.634093 | Cohesiveness |
| RT | 72h | WT | 0.668668 | 0.0269 | 40 | 0.579962 | 0.770942 | Cohesiveness |
| RT | 72h | *sbeII* | 0.676337 | 0.0272 | 40 | 0.586614 | 0.779784 | Cohesiveness |
| Fresh | 0h | WT | 733.626 | 68.894 | 40 | 526.3783 | 1022.472 | Cohesiveness |
| Fresh | 0h | *sbeII* | 515.8861 | 48.447 | 40 | 370.1494 | 719.0027 | Cohesiveness |
| Freezer | 24h | WT | 543.2563 | 51.017 | 40 | 389.7876 | 757.1492 | Cohesiveness |
| Freezer | 24h | *sbeII* | 468.7169 | 44.017 | 40 | 336.3055 | 653.2619 | Cohesiveness |
| Fridge | 24h | WT | 809.168 | 75.988 | 40 | 580.5799 | 1127.757 | Cohesiveness |
| Fridge | 24h | *sbeII* | 729.0972 | 68.469 | 40 | 523.1289 | 1016.16 | Cohesiveness |
| RT | 24h | WT | 609.2306 | 57.212 | 40 | 437.1243 | 849.0992 | Cohesiveness |
| RT | 24h | *sbeII* | 554.4715 | 52.07 | 40 | 397.8346 | 772.7801 | Cohesiveness |
| Freezer | 48h | WT | 524.144 | 47.791 | 35 | 378.3489 | 726.1204 | Cohesiveness |
| Freezer | 48h | *sbeII* | 457.1171 | 42.928 | 40 | 327.9825 | 637.0949 | Cohesiveness |
| Fridge | 48h | WT | 807.6437 | 75.845 | 40 | 579.4862 | 1125.632 | Cohesiveness |
| Fridge | 48h | *sbeII* | 681.4935 | 63.999 | 40 | 488.9731 | 949.8137 | Cohesiveness |
| RT | 48h | WT | 690.9314 | 64.885 | 40 | 495.7449 | 962.9676 | Cohesiveness |
| RT | 48h | *sbeII* | 601.9149 | 56.525 | 40 | 431.8753 | 838.9032 | Cohesiveness |
| Freezer | 72h | WT | 593.2733 | 55.714 | 40 | 425.6749 | 826.8591 | Cohesiveness |
| Freezer | 72h | *sbeII* | 503.3437 | 47.269 | 40 | 361.1502 | 701.5221 | Cohesiveness |
| Fridge | 72h | WT | 1088.233 | 102.2 | 40 | 780.8099 | 1516.697 | Cohesiveness |
| Fridge | 72h | *sbeII* | 634.5509 | 59.59 | 40 | 455.2917 | 884.3887 | Cohesiveness |
| RT | 72h | WT | 888.9669 | 83.482 | 40 | 637.8358 | 1238.974 | Cohesiveness |
| RT | 72h | *sbeII* | 593.9452 | 55.777 | 40 | 426.1571 | 827.7956 | Cohesiveness |

| | | | | | | | | |
|---|---|---|---|---|---|---|---|---|
| Fresh | 0h | WT | 0.642881 | 0.0144 | 40 | 0.593882 | 0.695922 | Cohesiveness |
| Fresh | 0h | *sbeII* | 0.626407 | 0.014 | 40 | 0.578664 | 0.67809 | Cohesiveness |
| Freezer | 24h | WT | 0.577142 | 0.0129 | 40 | 0.533153 | 0.62476 | Cohesiveness |
| Freezer | 24h | *sbeII* | 0.523458 | 0.0117 | 40 | 0.483561 | 0.566647 | Cohesiveness |
| Fridge | 24h | WT | 0.471252 | 0.0106 | 40 | 0.435334 | 0.510133 | Cohesiveness |
| Fridge | 24h | *sbeII* | 0.519163 | 0.0116 | 40 | 0.479593 | 0.561997 | Cohesiveness |
| RT | 24h | WT | 0.544125 | 0.0122 | 40 | 0.502653 | 0.589019 | Cohesiveness |
| RT | 24h | *sbeII* | 0.552746 | 0.0124 | 40 | 0.510617 | 0.598351 | Cohesiveness |
| Freezer | 48h | WT | 0.581027 | 0.0129 | 38 | 0.537069 | 0.628583 | Cohesiveness |
| Freezer | 48h | *sbeII* | 0.492143 | 0.011 | 40 | 0.454633 | 0.532747 | Cohesiveness |
| Fridge | 48h | WT | 0.435805 | 0.0098 | 40 | 0.402589 | 0.471762 | Cohesiveness |
| Fridge | 48h | *sbeII* | 0.469281 | 0.0105 | 40 | 0.433513 | 0.507999 | Cohesiveness |
| RT | 48h | WT | 0.489392 | 0.011 | 40 | 0.452092 | 0.52977 | Cohesiveness |
| RT | 48h | *sbeII* | 0.505461 | 0.0113 | 40 | 0.466936 | 0.547165 | Cohesiveness |
| Freezer | 72h | WT | 0.551776 | 0.0124 | 40 | 0.509721 | 0.597301 | Cohesiveness |
| Freezer | 72h | *sbeII* | 0.481038 | 0.0108 | 40 | 0.444374 | 0.520726 | Cohesiveness |
| Fridge | 72h | WT | 0.37733 | 0.0085 | 40 | 0.348571 | 0.408462 | Cohesiveness |
| Fridge | 72h | *sbeII* | 0.474302 | 0.0106 | 40 | 0.438152 | 0.513435 | Cohesiveness |
| RT | 72h | WT | 0.450442 | 0.0101 | 40 | 0.416111 | 0.487607 | Cohesiveness |
| RT | 72h | *sbeII* | 0.487987 | 0.0109 | 40 | 0.450794 | 0.528249 | Cohesiveness |
| Fresh | 0h | WT | 277.0349 | 26.774 | 40 | 196.8383 | 389.9055 | Chewiness |
| Fresh | 0h | *sbeII* | 320.864 | 31.01 | 40 | 227.9797 | 451.5915 | Chewiness |
| Freezer | 24h | WT | 474.6306 | 45.87 | 40 | 337.2337 | 668.0064 | Chewiness |
| Freezer | 24h | *sbeII* | 473.1832 | 45.73 | 40 | 336.2052 | 665.9692 | Chewiness |
| Fridge | 24h | WT | 812.071 | 78.482 | 40 | 576.9911 | 1142.928 | Chewiness |
| Fridge | 24h | *sbeII* | 670.8578 | 64.834 | 40 | 476.6566 | 944.1811 | Chewiness |
| RT | 24h | WT | 610.4644 | 58.998 | 40 | 433.746 | 859.1821 | Chewiness |
| RT | 24h | *sbeII* | 452.1936 | 43.702 | 40 | 321.2917 | 636.4279 | Chewiness |
| Freezer | 48h | WT | 460.4346 | 44.102 | 39 | 327.8144 | 646.7075 | Chewiness |
| Freezer | 48h | *sbeII* | 447.477 | 43.246 | 40 | 317.9405 | 629.7897 | Chewiness |
| Fridge | 48h | WT | 804.2747 | 77.728 | 40 | 571.4518 | 1131.955 | Chewiness |

| | | | | | | | | |
|---|---|---|---|---|---|---|---|---|
| Fridge | 48h | *sbeII* | 654.5011 | 63.254 | 40 | 465.0349 | 921.1603 | Chewiness |
| RT | 48h | WT | 691.7454 | 66.853 | 40 | 491.4977 | 973.5789 | Chewiness |
| RT | 48h | *sbeII* | 485.5551 | 46.926 | 40 | 344.9957 | 683.3818 | Chewiness |
| Freezer | 72h | WT | 590.1039 | 57.03 | 40 | 419.2795 | 830.5262 | Chewiness |
| Freezer | 72h | *sbeII* | 498.8946 | 48.215 | 40 | 354.4737 | 702.1561 | Chewiness |
| Fridge | 72h | WT | 1088.795 | 105.23 | 40 | 773.6088 | 1532.396 | Chewiness |
| Fridge | 72h | *sbeII* | 627.5834 | 60.652 | 40 | 445.9094 | 883.2757 | Chewiness |
| RT | 72h | WT | 886.9733 | 85.721 | 40 | 630.2106 | 1248.347 | Chewiness |
| RT | 72h | *sbeII* | 594.0192 | 57.408 | 40 | 422.0614 | 836.0366 | Chewiness |
| Fresh | 0h | WT | 2.90255 | 0.2297 | 42 | 2.195948 | 3.836519 | Resilience |
| Fresh | 0h | *sbeII* | 1.397544 | 0.1155 | 50 | 1.048093 | 1.863508 | Resilience |
| Freezer | 24h | WT | 1.144744 | 0.0888 | 39 | 0.869596 | 1.506951 | Resilience |
| Freezer | 24h | *sbeII* | 0.99051 | 0.0768 | 39 | 0.752434 | 1.303916 | Resilience |
| Fridge | 24h | WT | 0.996651 | 0.0773 | 39 | 0.757098 | 1.312 | Resilience |
| Fridge | 24h | *sbeII* | 1.086795 | 0.0843 | 39 | 0.825576 | 1.430666 | Resilience |
| RT | 24h | WT | 0.998034 | 0.0774 | 39 | 0.758149 | 1.31382 | Resilience |
| RT | 24h | *sbeII* | 1.016732 | 0.0822 | 45 | 0.765937 | 1.349645 | Resilience |
| Freezer | 48h | WT | 1.137755 | 0.087 | 37 | 0.866478 | 1.493963 | Resilience |
| Freezer | 48h | *sbeII* | 1.021416 | 0.0792 | 39 | 0.775911 | 1.344601 | Resilience |
| Fridge | 48h | WT | 1.004173 | 0.0779 | 39 | 0.762812 | 1.321902 | Resilience |
| Fridge | 48h | *sbeII* | 1.041348 | 0.0808 | 39 | 0.791052 | 1.370839 | Resilience |
| RT | 48h | WT | 0.998964 | 0.0775 | 39 | 0.758856 | 1.315045 | Resilience |
| RT | 48h | *sbeII* | 1.027492 | 0.0831 | 45 | 0.774043 | 1.363928 | Resilience |
| Freezer | 72h | WT | 1.005527 | 0.078 | 39 | 0.763841 | 1.323684 | Resilience |
| Freezer | 72h | *sbeII* | 1.00899 | 0.0783 | 39 | 0.766471 | 1.328243 | Resilience |
| Fridge | 72h | WT | 0.999493 | 0.0775 | 39 | 0.759257 | 1.315741 | Resilience |
| Fridge | 72h | *sbeII* | 1.011149 | 0.0784 | 39 | 0.768111 | 1.331085 | Resilience |
| RT | 72h | WT | 1.002361 | 0.0777 | 39 | 0.761436 | 1.319517 | Resilience |
| RT | 72h | *sbeII* | 0.999844 | 0.0775 | 39 | 0.759524 | 1.316203 | Resilience |
| Fresh | 0h | WT | 277.0349 | 26.774 | 40 | 196.8383 | 389.9055 | Gumminess |
| Fresh | 0h | *sbeII* | 320.864 | 31.01 | 40 | 227.9797 | 451.5915 | Gumminess |

| | | | | | | | | |
|---|---|---|---|---|---|---|---|---|
| Freezer | 24h | WT | 474.6306 | 45.87 | 40 | 337.2337 | 668.0064 | Gumminess |
| Freezer | 24h | *sbeII* | 473.1832 | 45.73 | 40 | 336.2052 | 665.9692 | Gumminess |
| Fridge | 24h | WT | 812.071 | 78.482 | 40 | 576.9911 | 1142.928 | Gumminess |
| Fridge | 24h | *sbeII* | 670.8578 | 64.834 | 40 | 476.6566 | 944.1811 | Gumminess |
| RT | 24h | WT | 610.4644 | 58.998 | 40 | 433.746 | 859.1821 | Gumminess |
| RT | 24h | *sbeII* | 452.1936 | 43.702 | 40 | 321.2917 | 636.4279 | Gumminess |
| Freezer | 48h | WT | 460.4346 | 44.102 | 39 | 327.8144 | 646.7075 | Gumminess |
| Freezer | 48h | *sbeII* | 447.477 | 43.246 | 40 | 317.9405 | 629.7897 | Gumminess |
| Fridge | 48h | WT | 804.2747 | 77.728 | 40 | 571.4518 | 1131.955 | Gumminess |
| Fridge | 48h | *sbeII* | 654.5011 | 63.254 | 40 | 465.0349 | 921.1603 | Gumminess |
| RT | 48h | WT | 691.7454 | 66.853 | 40 | 491.4977 | 973.5789 | Gumminess |
| RT | 48h | *sbeII* | 485.5551 | 46.926 | 40 | 344.9957 | 683.3818 | Gumminess |
| Freezer | 72h | WT | 590.1039 | 57.03 | 40 | 419.2795 | 830.5262 | Gumminess |
| Freezer | 72h | *sbeII* | 498.8946 | 48.215 | 40 | 354.4737 | 702.1561 | Gumminess |
| Fridge | 72h | WT | 1088.795 | 105.23 | 40 | 773.6088 | 1532.396 | Gumminess |
| Fridge | 72h | *sbeII* | 627.5834 | 60.652 | 40 | 445.9094 | 883.2757 | Gumminess |
| RT | 72h | WT | 886.9733 | 85.721 | 40 | 630.2106 | 1248.347 | Gumminess |
| RT | 72h | *sbeII* | 594.0192 | 57.408 | 40 | 422.0614 | 836.0366 | Gumminess |
| Fresh | 0h | WT | 2.90255 | 0.2297 | 42 | 2.195948 | 3.836519 | Springiness |
| Fresh | 0h | *sbeII* | 1.397544 | 0.1155 | 50 | 1.048093 | 1.863508 | Springiness |
| Freezer | 24h | WT | 1.144744 | 0.0888 | 39 | 0.869596 | 1.506951 | Springiness |
| Freezer | 24h | *sbeII* | 0.99051 | 0.0768 | 39 | 0.752434 | 1.303916 | Springiness |
| Fridge | 24h | WT | 0.996651 | 0.0773 | 39 | 0.757098 | 1.312 | Springiness |
| Fridge | 24h | *sbeII* | 1.086795 | 0.0843 | 39 | 0.825576 | 1.430666 | Springiness |
| RT | 24h | WT | 0.998034 | 0.0774 | 39 | 0.758149 | 1.31382 | Springiness |
| RT | 24h | *sbeII* | 1.016732 | 0.0822 | 45 | 0.765937 | 1.349645 | Springiness |
| Freezer | 48h | WT | 1.137755 | 0.087 | 37 | 0.866478 | 1.493963 | Springiness |
| Freezer | 48h | *sbeII* | 1.021416 | 0.0792 | 39 | 0.775911 | 1.344601 | Springiness |
| Fridge | 48h | WT | 1.004173 | 0.0779 | 39 | 0.762812 | 1.321902 | Springiness |
| Fridge | 48h | *sbeII* | 1.041348 | 0.0808 | 39 | 0.791052 | 1.370839 | Springiness |
| RT | 48h | WT | 0.998964 | 0.0775 | 39 | 0.758856 | 1.315045 | Springiness |

| | | | | | | | | |
|---|---|---|---|---|---|---|---|---|
| RT | 48h | *sbeII* | 1.027492 | 0.0831 | 45 | 0.774043 | 1.363928 | Springiness |
| Freezer | 72h | WT | 1.005527 | 0.078 | 39 | 0.763841 | 1.323684 | Springiness |
| Freezer | 72h | *sbeII* | 1.00899 | 0.0783 | 39 | 0.766471 | 1.328243 | Springiness |
| Fridge | 72h | WT | 0.999493 | 0.0775 | 39 | 0.759257 | 1.315741 | Springiness |
| Fridge | 72h | *sbeII* | 1.011149 | 0.0784 | 39 | 0.768111 | 1.331085 | Springiness |
| RT | 72h | WT | 1.002361 | 0.0777 | 39 | 0.761436 | 1.319517 | Springiness |
| RT | 72h | *sbeII* | 0.999844 | 0.0775 | 39 | 0.759524 | 1.316203 | Springiness |

*Table 5. Proportion of chain length distribution per DP fraction. Data is reported as Mean ± SD of n = 3, p-value obtained from independent samples t-test between sbeII and WT control starch*

| **Genotype** | **WT** | ***sbeII*** | ***p-value*** |
|---|---|---|---|
| 0-12 DP | 25.1 ± 2.7 | 14.1 ± 2.2 | 0.005 |
| 12-25 DP | 27.3 ± 1.5 | 17.5 ± 1.9 | 0.002 |
| 25-37 DP | 10.8 ± 0.3 | 9.6 ± 0.9 | 0.13 |
| 37-100 DP | 10.9 ± 0.7 | 19.0 ± 0.3 | 0.0006 |
| 100-1600 DP | 15.8 ± 2.7 | 26.3 ± 3.4 | 0.01 |
| >1600 DP | 10.1 ± 1.2 | 13.5 ± 1.7 | 0.05 |

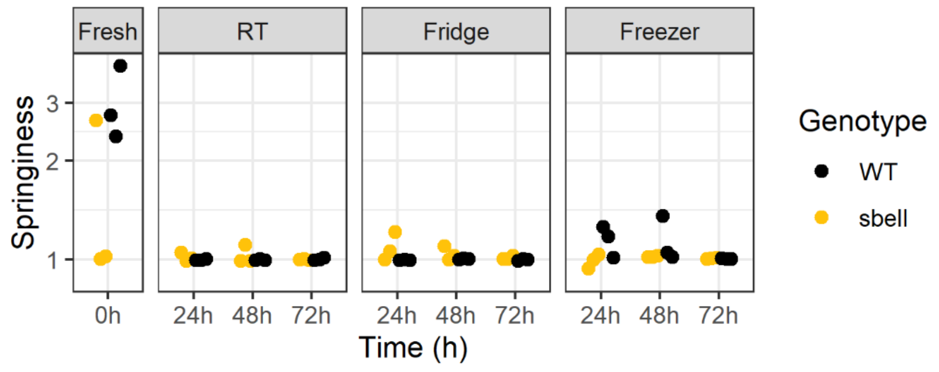


*Figure 1. Springiness of bread over storage obtained from texture analysis of bread crumb.*

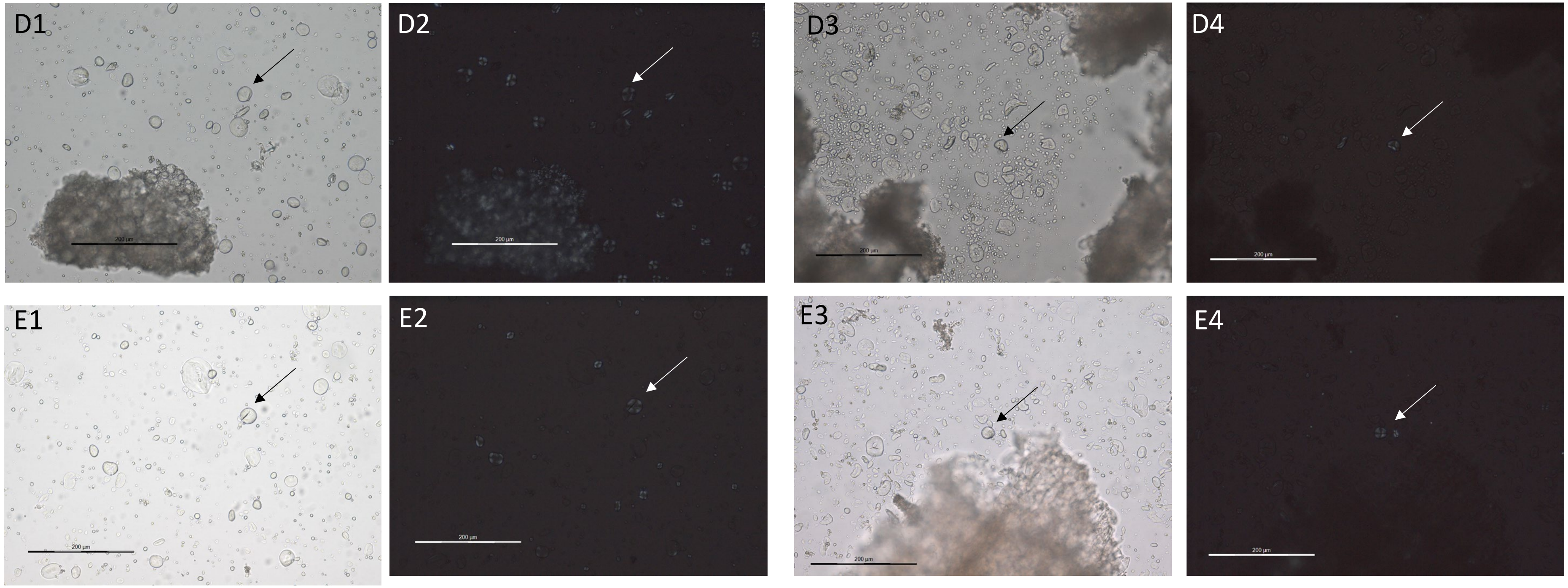


*Figure 2 Micrographs of starch in water using brightfield (BF) and polarised light filter (P) highlighting birefringence of starch granules. 1. WT control starch under BF; 2. WT control starch under P; 3. sbeII mutant starch under BF; 4. sbeII mutant starch under P; D. Starch from fresh bread crust; E: Starch from frozen crust after 7 days of freezer storage.*

Kweon, M., Slade, L., & Levine, H. (2011). Solvent Retention Capacity (SRC) Testing of Wheat Flour: Principles and Value in Predicting Flour Functionality in Different Wheat-Based Food Processes and in Wheat Breeding—A Review. *Cereal Chemistry, 88*(6), 537-552.
